# Rice *TSV3* Encoding Obg-like GTPase Protein is Essential for Chloroplast Development during the Early Leaf Stage under Cold Stress

**DOI:** 10.1101/163576

**Authors:** Dongzhi Lin, Quan Jiang, Xiaojing Ma, Kailun Zheng, Xiaodi Gong, Sheng Teng, Jianlong Xu, Yanjun Dong

## Abstract

The Spo0B-associated GTP-binding (Obg) proteins occupy a wide variety of roles in the viability of nearly all bacteria. Its detailed roles in higher plants have not yet been elucidated. A novel rice thermo-sensitive virescent mutant *tsv3* was identified in this study that displayed albino phenotype at 20°C before the 3-leaf stage while being normal green at 32°C or even at 20°C after the 4-leaf stage. The mutant phenotype was aligned with altered chlorophyll (Chl) content and chloroplast development. Map-based cloning and complementation test showed that *TSV3* encoded a kind of small GTP binding protein. Subcellular localization revealed that TSV3 was in chloroplast. *TSV3* transcripts were highly expressed in leaves and weak or undetectable in other tissues, suggesting the tissue-specific expression. In *tsv3* mutant, the transcriptional levels of certain genes associated with biogenesis of chloroplast ribosomes 50S subunit were severely decreased at the 3-leaf-stage under cold stress, but could be recovered to normal levels at a higher temperature (32°C). The observations from this study indicated that the rice nuclear-encoded *TSV3* plays important roles in chloroplast development at early leaf stage under cold stress.

---

Chloroplast is a semi-autonomous organelle containing many genes important for metabolic pathways of photosynthesis (Mandel et al. 1996). Chloroplast development during leaf development consists of a series of complex events associated with chloroplast differentiation and can be divided into three steps coordinately regulated by the plastid and nucleus genes (Mullet 1993; Kusumi et al. 2011). The first step involves the activation of plastid replication and plastid DNA synthesis. The second step is the chloroplast “build-up”, characterized by the establishment of chloroplast genetic system. At this step nuclear-encoded plastid RNA polymerase (NEP) preferentially transcribes plastid genes encoding plastid gene expression machineries (Hajdukiewicz et al.1997), and the transcription and translation activities in the chloroplast are dramatically increased. At third step, the plastid and nuclear genes encoding photosynthetic apparatus are expressed at very high levels. Plastid genes for photosynthetic apparatus are predominantly transcribed by plastid-encoded RNA polymerase (PEP) (Santis-Maciossek et al., 1999). All expressions of these genes lead to the synthesis and assembly of chloroplast. In spite of these, the mechanisms of the major genes in higher plants remain largely unknown (Pfalz and Pfannschmidt 2012).

GTPases are a large family of enzymes that hydrolyze guanosine triphosphate (GTP). GTPases are uncovered universally in all kingdoms of life and play a crucial role in many cellular processes (Bourne et al. 1990). The Spo0B-associated GTP-binding (Obg) protein subfamily of GTPases was originally identified at downstream of the Spo0B gene in *Bacillus subtilis* (Trach and Hoch 1989). Typical Obgs are large GTPases which contained three domains, i.e., the Obg fold, G domain and Obg C-terminal region (OCT) (Buglino et al. 2002; Kukimoto-Niino et al. 2004). The Obgs proteins have been shown to be essential for the viability of nearly all bacteria (Maddock et al. 1997; Okamoto and Ochi 1998; Shepherd et al. 2002; Foti et al. 2005; Michel 2005). It is noteworthy that the majority of Obg proteins studied to date are associated with the ribosome except that ObgE is involved in *Escherichia coli* chromosome partitioning, partially associated with the membrane (Kobayashi et al. 2001). For example, Obgs from *Bacillus subtilis, Caulobacter crescentus* and *Escherichia coli* have been reported to be associated with the 50S ribosomal subunit (Scott et al. 2000; Lin et al. 2004; Wout et al. 2004) and the mutations have been shown to affect ribosome assembly or maturation (Datta et al. 2004; Jiang et al. 2006). Recently, two mutations in Obg fold and OCT region impaired the ability of *Bacillus* Obg to associate with ribosomes and to induce a general stress regulation, speculating that Obg may have dual functions in ribosome biogenesis and stress responses (Kuo et al. 2008). In eukaryotic cell, certain studies showed that an *Arabidopsis* Obg homolog, AtObgC/CPSAR1, localized both the inner envelope and the stroma of chloroplasts, is essential for the formation of normal thylakoid membranes (Garcia et al. 2010); However, others reported it may play an important role in the biogenesis of chloroplast ribosomes (Bang et al. 2009). More recently, based on results of studies on *Arabidopsis AtObgC/CPSAR1* and rice *OsObgC*, Bang et al. (2012) reported that ObgC functions primarily in plastid ribosome biogenesis during chloroplast development.

Rice (*Oryza sativa*) mutants are ideal materials to explicate the function of chloroplast development in higher plants. In this study, a new rice thermo-sensitive virescent mutant *tsv3*, which exhibited the albino phenotype before the 3-leaf stage at 20°C and normal green at 32°C or even at 20°C after the 4-leaf stage, was used. The mutation of rice *TSV3*, encoding the Obg subfamily of small GTP-binding protein, was responsible for the mutant phenotype. Additionally, the transcript levels of genes associated with chlorophyll biosynthesis and photosynthesis, of some genes associated with biogenesis of chloroplast ribosomes 50S subunit were severely affected in *tsv3* mutants at low temperature. The findings implicated rice *TSV3* plays an important role in chloroplast development during leaf development under cold stress.

## MATERIALS and METHODS

### Plant materials and growth conditions

The rice thermo-sensitive virescent mutant *tsv3* was discovered in our mutant pool from Jiahua 1 (WT, *japonica* rice variety) irradiated with ^60^Co gamma rays in 2006. The F_2_ population for genetic mapping was generated from a cross between Pei’ai 64S (*indica*) and *tsv3* mutant. The thermo-sensitive virescent phenotype in *tsv3* mutant can be distinguished from normal green at Hainan (winter season, subtropical climate) and Shanghai (spring season, temperate climate), China under local growing conditions. WT and *tsv3* plants were grown in growth chambers under controlled 12 h of light and 12 h of dark at a constant temperature of 20°C and 32°C, respectively, for phenotypic characterization, pigment content measurement and RNA extraction.

### Chlorophyll (Chl) and carotenoid (Car) content measurement

Both Chl and Car contents were assessed using a spectrophotometer following the slightly modified methods of Arnon (1949) and Wellburn (1994). Briefly, fresh leaves (0.2 g each sample) from the 3-leaf-stage seedlings grown at 20°C and 32°C, respectively, were fetched, cut and homogenized in 5 mL of acetone:ethanol:H_2_O (by 5:4:1 vol.) for 18 h under dark conditions, progressively. Residual debris was removed by centrifugation. The supernatants were analyzed with a UV5100 Spectrophotometer (Beckman Coulter, USA) at 663, 645 and 470 nm.

### Transmission electron microscopy (TEM)

For TEM analysis, transverse sections were sampled from the same parts of the top leaves at the 3-leaf-stage of seedlings grown at 20°C and 32°C, respectively. Samples were fixed in a solution of 2.5% glutaraldehyde first, then in 1% OsO_4_ buffer at 4°C for 5h after vacuum. After staining with uranyl acetate, tissues were further dehydrated in an ethanol series and finally embedded in Spurr’s medium prior to ultrathin sectioning. Samples were stained again and examined with a Hitachi-7650 (Hitachi, Tokyo, Japan) transmission electron microscope.

### Mapping of *TSV3* gene

Rice genomic DNA was extracted from fresh leaves by the modified CTAB method (Murray and Thompson 1980). Totally, 1,430 plants with the mutant phenotype were selected from F_2_ populations for mapping of the *TSV3* locus. Initially, we adopted 81 SSR primers based on the Gramene database (http://www.gramene.org). New SSR and InDel markers were developed based on the entire genomic sequences of the *japonica* Nipponbare variety (Goff et al 2002) and the *indica* variety 9311 (Yu et al. 2002). Details of the markers used for mapping were listed in Supplemental Table 1. The genomic DNA fragments of the candidate genes from the mutant and WT plants were amplified and sequenced. The sequencing reaction was performed by Sinogenomax Co., Ltd. The function and ORFs of the candidate genes were obtained from TIGR (http://rice.plantbiology.msu.edu/cgi-bin/gbrowse/rice/). Conserved domain structures were predicted through SMART (http://smart.embl-heidelberg.de/).

### RT-PCR and realtime PCR (qPCR) analysis

To examine the expression pattern of *TSV3*, the WT RNA was extracted from germinating bud, plumules, roots, stems and leaves at seedling-stage, flag leaves and young panicles at heading stage using an RNA Prep Pure Plant kit (Tiangen Co., Beijing, China) and was reversely transcribed by ReverTra Ace (ToYoBo, Osaka, Japan) following the manufacturer’s instructions. RT-PCR analysis was carried out to assess *TSV3* transcript levels. For transcriptional analysis of *TSV3* and other 23 genes associated with Chl biosynthesis, chloroplast development and photosynthesis (*HEMA1, CAO1, YGL1, PORA, Cab1R, RbcS, RbcL, PsaA, PsbA, LhcpII, RNRS, RNRL, V2, OsRpoTp, OsPoLP1, FtsZ, RpoA, RpoB, Rps7, Rps20, Rpl21, 16SrRNA* and *23SrRNA*) in rice, total RNA was extracted from the 3^rd^ leaves of WT and *tsv3* plants. qPCR analyses were performed using a SYBR Premix Ex TaqTM kit (TaKaRa) on an ABI7500 Realtime PCR System (Applied Biosystems; http://www.appliedbiosystems.com), and the relative quantification of gene expression data was performed as described by Livak and Schmittgen (2001). The specific primers for qPCR were designed according to both Wu et al. (2007) and NCBI-published sequences and were listed in Supplemental Table 2. The rice *Actin* gene was used as a reference gene.

### Complementation test

A 8.3-kb genomic DNA fragment covering the entire *TSV3* gene, plus each 2.0 kb upstream and downstream sequence, was amplified from the WT parent with the primer pair *TSV3* F: 5'-GGGGTACCCCTTGACATACCTCTCCTGTTTGC-3' and *TSV3* R:5'-CGGGATCCCGCTGGGTTGGACAGATAATATGC-3'. The underlined sequences represent the cleavage sites of *Kpn*I and *BamH*I, respectively. The PCR product was ligased with the pMD18-T vector (TaKaRa, Japan), and the fragment was subcloned into the pCAMBIA1301 binary vector (CAMBIA; http://www.cambia.org.au) after sequence verification. The resultant pCAMBIA1301-TSV3 plasmid and the empty vector as control were transferred into *Agrobacterium tumefaciens* EHA105 and were introduced into the *tsv3* mutant via agrobacterium-mediated transformation (Hiei et al. 1994). The genotype of transgenic plants was determined by PCR amplification of the *hygromycin phosphotransferase* gene (*hpt*) with primers *HPTF* (5’-GGAGCATATACGCCCGGAGT-3’) and *HPTR*

(5’-GTTTATCGGCACTTTGCATCG-3’) and *GUS* gene with primers *GUSF* (5’-GGGATCCATCGCAGCGTAATG-3’) and *GUSR* (5’-GCCGACAGCAGCAGTTTCATC-3’) as selection.

### Subcellular localization

To investigate the subcellular localization of TSV3 protein, a cDNA fragment containing the N-terminal region (amino acids 1-280) of *TSV3* was amplified from total RNA in WT plants using primer pair 5'-GAAGATCTATGCCGCTCCTCCTCCAC-3' (restriction site of *BglII*); 5'-GGGGTACCCCACCAACGTCTGCAACCAC-3' (restriction site of *KpnI*) and was introduced into vector pMON530-GFP. The pMON530:CaMV35S:TSV3-GFP plasmid was then transferred into *Agrobacterium* EHA105 after sequence verification. For subcellular localization of TSV3, transient expression assays were performed in tobacco (*Nicotiana tabacum*) according to the method described by Jiang et al. (2014). The GFP fluorescence images were obtained using argon ion laser excitation of 488 nm with a 505–530 nm band-pass filter.

### Sequence and phylogenetic analyses

Gene prediction was performed using the Rice Genome Annotation Project database (RGAP, http://rice.plantbiology.msu.edu/). The full-length amino acid sequence of *TSV3* and most similar sequences identified via BLAST search were aligned with the MUSCLE tool (Edgar 2004) using the default parameters. A neighbor-joining tree was constructed using MEGA v5.2 software (http://www.megasoftware.net/; Tamura et al., 2011) by the bootstrap method with 1,000 replicates. Multiple sequence alignments were conducted with BioEdit software (http://www.mbio.ncsu.edu/BioEdit/bioedit.html; Hall 1999).

## RESULTS

### Phenotype characterization of the tsv3 mutant

The leaves of *tsv3* mutant only appeared albino phenotype before the 4-leaf stage when grown at 20°C (Figure 1A,C), gradually turned yellowish green, and finally normal green after the 4-leaf stage even at 20°C or a higher temperature (32°C) (Figure 1B). Consistent with the mutant phenotype, the Chl a, Chl b, and Car contents in *tsv3* 3^rd^ leaves were drastically lower than those in WT at 20°C (Figure 1D); however, they were comparable between WT and *tsv3* plants at 32°C (Figure 1E). The observations showed that the *tsv3* mutant has the thermo-sensitivity of virescent phenotype at the early seedling stage.

**Figure 1.**
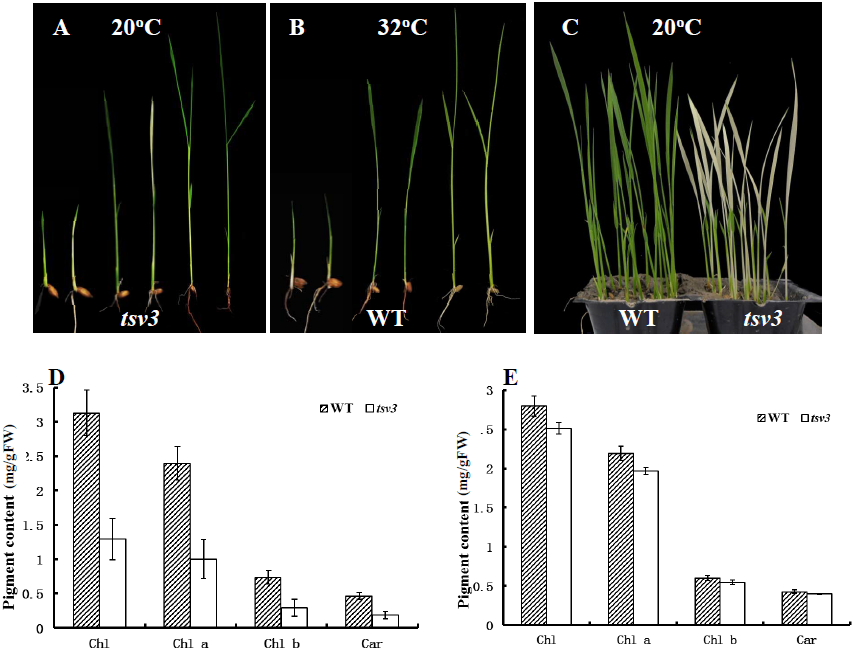
Characterization of the *tsv3* mutants. A 2-, 3- and 4-leaf stage plants of wild type (WT, Jiahua1, *left*) and *tsv3* mutant (*right*) at 20°C. B 2-, 3- and 4-leaf stage plants of WT (*left*) and *tsv3* mutant (*right*) at 32°C. C 3-leaf-stage-seedlings for WT(*left*) and *tsv3* mutant(*right*) grown at 20°C, respectively. D, E The pigment contents of the 3^rd^ leaves at 3-leaf stage at 20°C and 32°C for WT and *tsv3* mutant.

TEM analysis of chloroplasts was performed to examine if the lack of photosynthetic pigments in the *tsv3* mutant at low temperatures was accompanied by chloroplast ultrastructural changes. As it was expected, the grana lamella stacks in WT plants were dense and well-structured, regardless of lower or higher temperatures (Figure 2A, C). In *tsv3* mutant, the chloroplast structure at 20°C was not intact and less grana lamella stacks (Figure 2D), however it exhibited well-developed lamella structures (Figure 2B) at 32°C similar to WT (Figure 2A,C), suggesting the *tsv3* mutation only resulted in malformed chloroplast before 4-leaf stage under cold stress (20°C). Moreover, except for slight reduction in plant-height after transplanting (Supplemental Figure 1A), certain yield-related traits such as, panicle number, grains per panicle and 1000-grain weight (Supplemental Figure 1B) had no significant differences between *tsv3* and WT plants, indicating that *tsv3* mutation did not have obviously negative effects on growth under the field condition.

**Figure 2.**
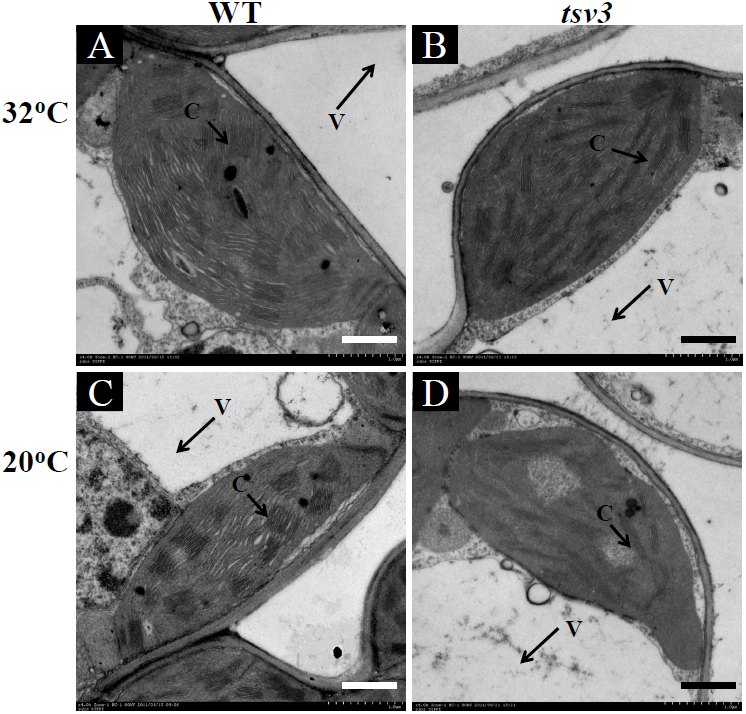
Transmission electron microscopic images of chloroplasts in WT and *tsv3* mutant at 20°C and 32°C. A, Intact chloroplast in the WT cell at 32°C. B Intact chloroplast in the *tsv3* mutant cell at 32°C. C Intact chloroplast in the WT cell at 20°C. D Abnormal chloroplast in the *tsv3* mutant cell at 20°C. G, grana stacks; V, vacuole

### Cloning of the *TSV3* gene

To understand the molecular mechanism responsible for the mutant phenotype, map-based cloning was used to identify the *TSV3* locus. Resultantly, the F_1_ plants from the cross of Pei’ai64S/*tsv3* were all normal green and the F_2_ population displayed a segregation pattern fitted a ratio of 3 to 1 (green: albino phenotype =453:132; χ^2^=3.45; *P* >0.05), demonstrating the albino phenotype is a recessive trait controlled by a single gene (*tsv3*). Initially, *tsv3* was mapped between the markers P1 and RM570 on chromosome 3 using 214 mutant individuals (Figure 3A). Ultimately, *tsv3* was narrowed to a 36kb interval between markers P2 and P5 on BAC clone AP104321 and no recombinant was found near the marker P3 and P4 (Figure 3A). Within this target region, seven candidate genes were predicted using the program RGAP (http://rice.plantbiology.msu.edu). All candidate genes were then sequenced and verified only four discontinuous nucleotide deletion (CAA^*^G) in the *tsv3* mutant (Figure 3A), at the first exon of *LOC_Os03g58540*, encoding an Obg subfamily of small GTP-binding protein, caused a premature stop codon of translation and resulted in a frame-shift mutation. In addition, the significant up-regulation for *LOC_Os03g58540* transcript in *tsv3* mutant, as compared with the WT plants at 20°C (Figure 3B), also indicated the possible existence of the *LOC_Os03g58540* mutation in *tsv3* mutant.

**Figure 3.**
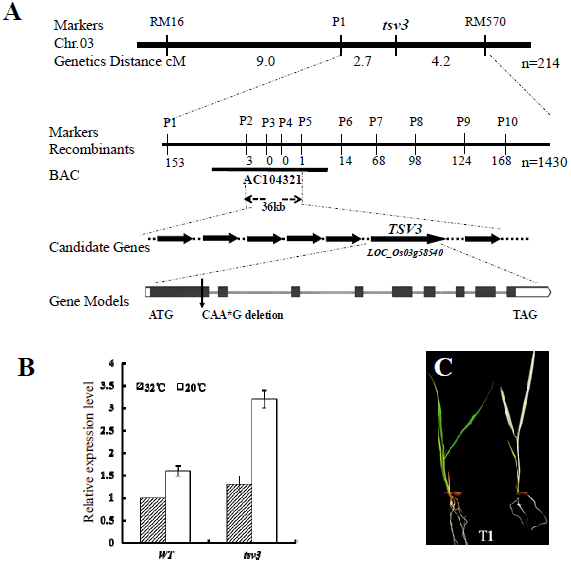
Genetic analysis and cloning of the *TSV3* gene. A Genetic mapping of the *TSV3* gene. B Transcript levels of *TSV3* (*LOC_Os03g58540*) in mutant and WT at the 3-leaf stage under different temperatures by qPCR. C The T_1_ segregation from transgenic T_0_ plants transformed with pCAMBIA1301-TSV3 at 20°C. The genotypes of green phenotype are *TSV3:TSV3* /*TSV3: tsv3* (*left*) and the genotype of the albino phenotype is *tsv3:tsv3* (*right*). OsActin was used as a control for qPCR.

To further confirm that the *LOC_Os03g58540* mutation was responsible for the mutant phenotype, a genetic complementation test was performed. Resultantly,all of the 23independent transgenic plants transformed with the vector of pCAMBIA1301:LOC_Os03g58540 driven by its own promoter were completely reverted to green leaf as WT plants. And the T_1_ generations derived from the transgenic T_0_ plants appeared the separation at 20°C (Figure 3C). This affirmed the *LOC_Os03g58540* corresponded to the *TSV3* gene.

### Characterization of TSV3 protein

TSV3 was predicted to be a 504-amino acid polypeptide with a calculated molecular mass of 54.5 kD. Conserved domain analysis showed that TSV3 contains a Spo0B-associated GTP-binding protein (the Obg subfamily of small GTP binding protein) and a 50S ribosomal subunit binding site domain by pfam (http://pfam.janelia.org).

Orthologs of TSV3 was found in *Arabidopsis thaliana, Populus trichocarpa, Ricinus communis, Vitis vinifera, Brachypodium, Glycine max, Sorghum bicolor and Zea mays* through NCBI and can be divided into two (I, II) groups (Figure 4B). Furthermore, group I can clearly be divided into two sub-branch(Ia,b). The sequences were highly conserved within higher plants, and TSV3 exhibited a maximum 84.9 % amino acid identity with orthologs TSV3 from *Zea mays* and shared 83.3 % similarity with orthologs TSV3 from *Sorghum bicolor* (Figure 4A). Notably, it was found the existence of three rice homologus Obg genes, *TSV3*, formerly termed as *OsObgC2* (Bang et al. 2009), *OsObgC1(LOC_Os07g47300*, Bang et al., 2009 and 2012), and *OsObgM* (*LOC_Os 11g47800*, Bang et al. 2009). As shown in Figure 4B, the TSV3 can be clearly divided into monocots and dicotyledons within Ib sub-branch. In addition, the predicted 3D structure (Supplemental Figure 2) of TSV3 (Ib branch) more likes both OsObgM and AtObgM (II group) than OsObgC1 (Ia branch).

**Figure 4.**
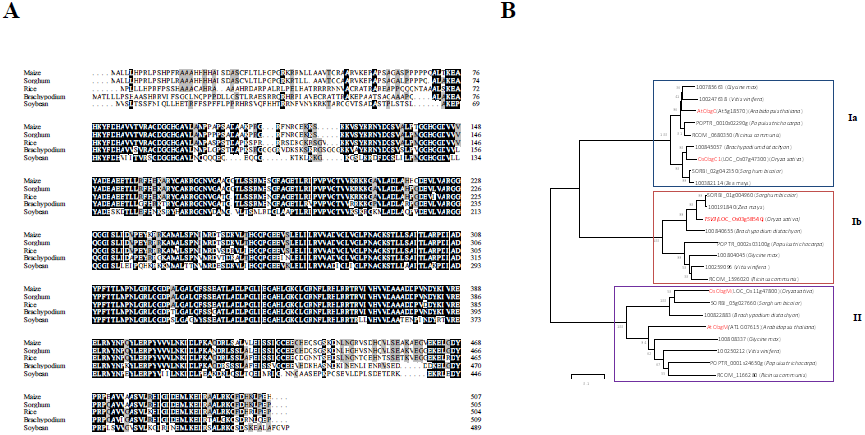
Phylogenic analysis of TSV3. A Amino acid sequence alignment of TSV3 with the four homologous proteins from Amino acids fully or partially conserved are shaded black and gray, respectively. B Phylogenic tree of TSV3 and homologous proteins. The rooted tree is based on a multiple sequence alignment generated with the program Mega5.2. Scale represents percentage substitution per site. Statistical support for the nodes is indicated.

### Subcellular localization of TSV3

The TSV3 was predicted to be localized in chloroplasts by TargetP (Emanuelsson et al. 2000, http://www.cbs.dtu.dk/services/TargetP/) and iPSORT (http://ipsort.hgc.jp/). To verify this prediction, the pMON530:CaMV35S:TSV3-GFP plasmid was introduced into tobacco cells for a transient expression assay. At the same time, the only GFP driven by the 35S promoter was transformed into tobacco cells as a control. Resultantly, in tobacco mesophyll cells transformed with the pMON530:CaMV35S:TSV3-GFP plasmid, GFP fluorescence perfectly overlapped with chloroplast autofluorescence (Figure 5B), while the empty GFP vector without a specific targeting sequence had green fluorescent signals in both the in cytoplasm and nucleus (Figure 5A). The results confirmed that TSV3 is localized in the chloroplast.

**Figure 5.**
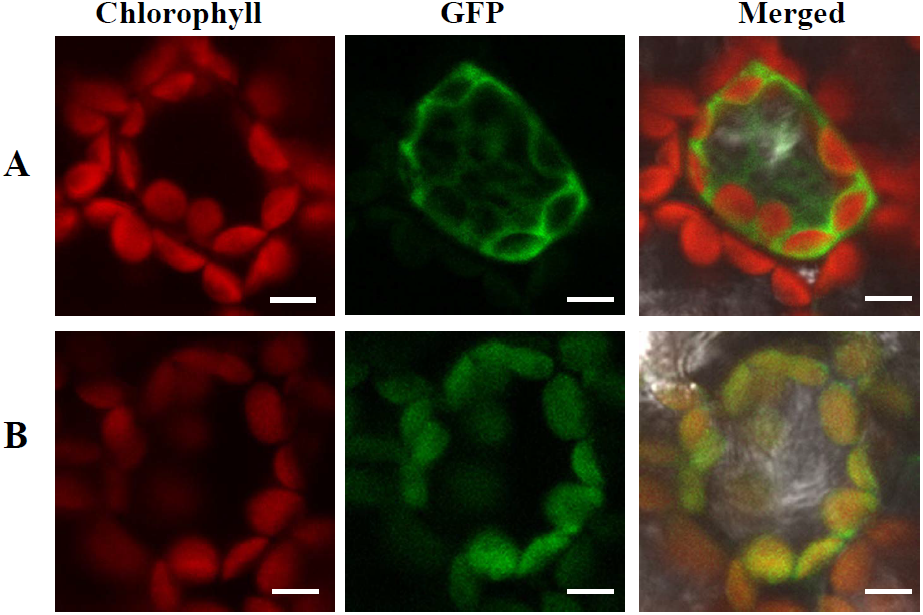
Subcellular localization of TSV3 protein. A Empty GFP vector without a specific targeting sequence. B TSV3-GFP fusion. The scale bar represents 20 μm.

### Expression pattern of *TSV3*

Reverse transcription (RT)-PCR was performed to examine the expression pattern of *TSV3*. A significantly high level of expression in seedling- and flag-leaves was detected, while weak expression in both roots and stems, and undetectable in both germinating bud and young panicles (Figure 6A), indicating that *TSV3* mainly functions in leaves. This was consistent with the data from rice gene expression profiling in the RiceXPro database (Supplemental Figure 3). In addition, the expression level of *TSV3* basically increased along with the leaf development from plumule to the 5th leaves (Figure 6B). It was noted that the *TSV3* transcript accumulation in the seedling-leaves was much more than that in the flag-leaves (Figure 6A). The results showed that *TSV3* might play an important role in leaf chloroplast development, especially for seedling stage.

**Figure 6.**
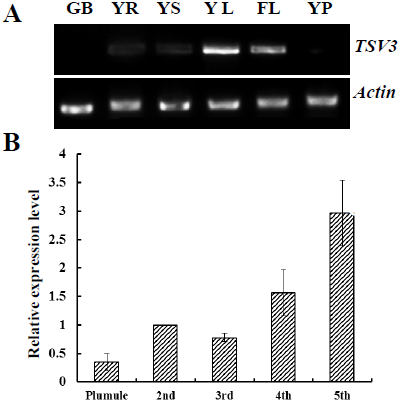
Expression analysis of *TSV3* by RT-PCR analysis. A Analysis of expression of *TSV3* in different tissues by RT-PCR. GB, Germinating bud; YR, young-seedling roots; YS, young-seedling stem; YL, young-seedling leaf; FL, flag leaf at heading; YP, young panicles. *OsActin* was used as a control (cycle number for *OsActin* was 28, cycle number for *TSV3* was 35). B Transcript levels of *TSV3* in top leaves sampled from the plumule, 2-, 3-, 4- and 5-leaf stages. The *TSV3* transcript level in the top leaves at the 2-leaf stage was set to 1.0, and the relative values in other treatments were calculated accordingly. *OsActin* was used as a control (Cycle number of *OsActin* is 28, cycle number of TSV3 is 31).

### The mutation of *TSV3* affects expression of associated genes

The transcript levels of genes for Chl biosynthesis, photosynthesis and chloroplast development both in the *tsv3* mutant and WT at 20°C and 32°C were examined. The expressions of genes for Chl biosynthesis (Wu et al. 2007), such as *CAO1* (*CHLOROPHYLLIDE A OXYGENASE1*), *HEMA1* (encoding glutamyl tRNA reductase), *YGL1* (encoding a Chl synthetase) and *PORA* (encoding NADPH-dependent protochlorophyllide oxidoreductase) were obviously down-regulated in the mutant (Figure 7A) at 20°C, in consistent with the decreased Chl content (Figure 1D) and the albino phenotype (Figure 1A, C). The photosynthesis associated transcripts from plastid genes *Cab1R* (encoding the light harvesting Chla/b-binding protein of PSII), *PsaA* and *PsbA* (encoding two reaction center polypeptides), and *RbcL* (encoding the large subunit of Rubisco) and the nuclear *RbcS* (encoding the small subunit of Rubisco) (Kyozuka et al. 1993) were greatly suppressed in the mutant at 20°C (Figure 7B). However, at 32°C, nearly all of transcriptional levels of the affected genes abovementioned were recovered to WT levels or slightly higher levels (Figure 8A, B).

**Figure 7.**
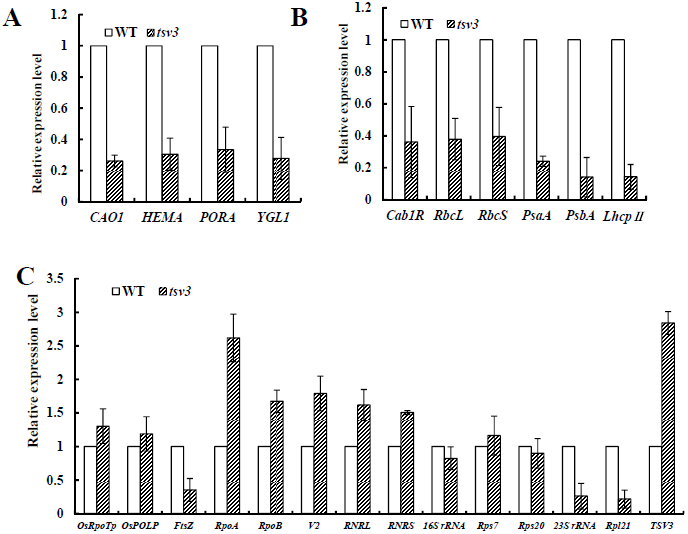
qPCR analysis of those genes related to Chl biosynthesis, photosynthesis and chloroplast development in mutant at 20°C. A, B, C Expression levels of genes related to Chl biosynthesis, photosynthesis and chloroplast development in WT and the *tsv3* mutant in the 3^rd^ leaves, respectively. The relative expression level of each gene in WT and mutant was analyzed by qPCR and normalized using the *OsActin* as an internal control. Data are means±SD (n = 3).

**Figure 8.**
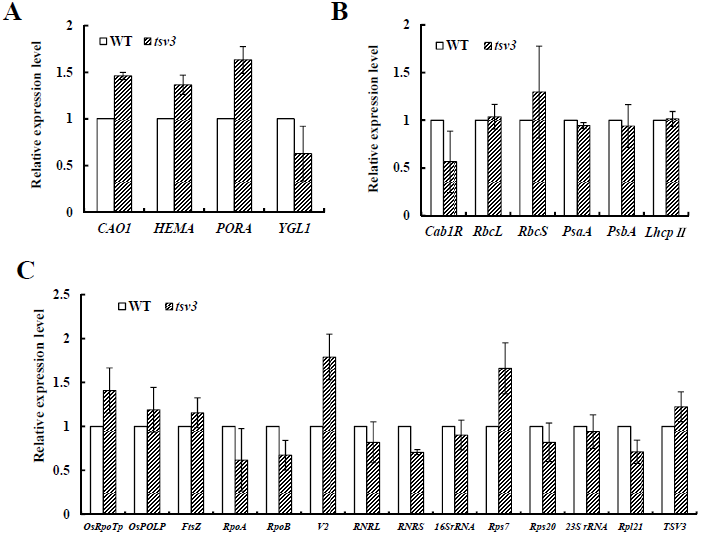
qPCR analysis of those genes related to Chl biosynthesis, photosynthesis and chloroplast development in mutant at 32°C. A, B, C Expression levels of genes related to Chl biosynthesis, photosynthesis and chloroplast development in WT and the *tsv3* mutant in the 3^rd^ leaves, respectively. The relative expression level of each gene in WT and mutant was analyzed by qPCR and normalized using the *OsActin* as an internal control. Data are means±SD (n = 3).

With regarding chloroplast-development associated genes, we investigated both nuclear-encoded genes [*RNRS* and *RNRL*, encoding the large and small subunits of ribonucleotide reductase (Yoo et al., 2009), *V2* encoding plastidal guanylate kinase (Sugimoto et al., 2007), *OsRpoTp* encoding NEP core subunits (Hiratsuka et al., 1989), *OsPoLP1* encoding one plastidial DNA polymerase (Vitha et al., 2001), *Rpl21* encoding the ribosomal protein L21 and *FtsZ* encoding a component of the plastid division machinery (Takeuchi et al., 2007)] and plastid-encoded genes [23S rRNA (23S ribosomal RNA), 16S rRNA (16S ribosomal RNA), *Rps7* encoding one ribosomal protein (Kusumi et al. 2011), *Rps20* encoding ribosomal protein S20, *RpoA* and *RpoB* encoding the PEP core α, β subunit (Kusumi et al. 2011)]. Resultantly, in the *tsv3* mutant, the transcriptional levels of *FtsZ, 23S rRNA* and *Rpl21* were remarkably reduced while other genes displayed WT levels or slightly higher levels at 20°C (Figure 7C). Interestingly, all affected genes were recovered to WT levels at 32°C (Figure 8C). It was possible that the abnormal expression of these key genes (*FtsZ, 23S rRNA* and *Rpl21*) led to the mutant phenotype under cold stress.

## DISCUSSION

In this study, we isolated a rice novel Obg subfamily of small GTP-binding protein gene *TSV3* by a map-based cloning strategy with a new rice thermo-sensitive virescent mutant *tsv3.* The virescent phenotype resulted from the four discrete deletion in *tsv3* mutant, which drastically affected the expression levels of some genes associated with Chl biosynthesis and photosynthesis (Figure 7A, B) and some key genes associated with chloroplast development such as *FtsZ, Rpl21* and *23SrRNA* (Figure 7C). This study demonstrates that the rice *TSV3* might play important role for early chloroplast development under cold stress.

### *TSV3* is needed for early chloroplast development under cold stress

At low temperature (20°C), the *tsv3* mutant seriously affected expressions of some genes associated with Chl biosynthesis, photosynthesis and chloroplast development (Figure 7A,B,C), whereas the expressions of those affected genes could back to normal level or slightly higher levels as WT plants at high temperature (32°C) (Figure 8A,B,C). This discrepancy was attributable to the difference in the chloroplast structure and pigment contents between 20°C and 32°C. It is clear that malfunction of *TSV3* influences early chloroplast development under cold stress.

Notwithstanding the reason why abnormal chloroplast occurs only in early leaves of rice under cold stress is not completely claimed yet, it was speculated that the *TSV3* function is possibly not prerequisite at higher temperatures, but it is essential/more required for rice chloroplast development during early-leaf development under cold stress. This was plausibly supported by the results from transcriptional analysis (Figure 3B) of the highly expressed level at 20°C, irrespective of WT or *tsv3* plants, and the no discrimination between WT and *tsv3* mutant at 32°C. Additionally, under cold stress (20°C), the decreased level of *FtsZ*, known to involve in the first step of chloroplast development, might affect the plastid division, which was in aligned with chloroplast development (data not shown) in mutant.

### The *TSV3* may regulate biogenesis of chloroplast 50S large ribosomal under cold stress

The chloroplast 50S large subunit consists of three rRNAs (23S, 4.5S, and 5S rRNAs) and 30S ribosomal proteins, e.g RPL21. Although much work has been undertaken to elucidate the composition of chloroplast ribosome, the molecular basis of its assembly in higher plants remains elusive. Previous studies on bacteria showed that the majority of Obg propteins are associated with the 50S ribosomal subunit, ribosome assembly and stress responses (Scott et al. 2000; Lin et al. 2004; Wout et al. 2004; Datta et al. 2004; Jiang et al. 2006; Kuo et al. 2008). In view of the previous results aforementioned, Obgs homolog *TSV3* possibly have similar functions in higher plants. Except for OCT region, the TSV3 homolog in *Arabidopsis*, namely AtObgC/CPSAR1 (At5g18570), and OsObgC1 (LOC_Os07g47300) in rice, share the similarity in 3D structure (Supplemental Figure 2). AtObgC/CPSAR1 was essential for the formation of normal thylakoid membranes (Garcia et al., 2010) and might play an important role in the biogenesis of chloroplast ribosomes (Bang et al., 2009). More recently, based on these results from previous studies on rice *OsObgC1* (*LOC_Os07g47300*) and *Arabidopsis AtObgC* (*At5g18570*), Bang et al. (2012) reported that plant *ObgC* is light-induced gene and its protein is translocated into chloroplast and then may be involved in biogenesis of chloroplast large (50S) ribosomal subunit, which influences the PEP-related plastid gene transcription, and proposed a hypothetical model of three ObgC domains (OCT, Obg fold and G domain), which may mediate the role ObgC of chloroplast ppGpp signaling, association of ObgC with 50S ribosomal subunit and may regulate the action of ObgC depending on its GTP-/GDP-bound states, respectively(Supplemental Figure 2).

In this study, the transcripts of *Rpl21* and *23SrRNA*, encoding the components of chloroplast ribosomal large subunit (50S), involved in the ribosome assembly, in the *tsv3* mutant were seriously affected under 20°C (Figure 7C), but those for *Rps7, Rps20* and *16SrRNA*, all encoding the components of chloroplast ribosomal small (30S) subunit, seemed to be no significant change at 20°C and 32°C (Figure 7G, Figure 8C). Additionally, the severely reduced transcription levels of PEP-dependent plastid genes (*RbcL, psaA, psbA*) (Figure 7B) suggested that the *tsv3* mutation affected the plastid-encoded RNA polymerase transcription, like A*tObgC* (Bang et al. 2012). Hence, like *AtObgC* (Bang et al. 2012), *TSV3* might be involved in biogenesis of chloroplast large (50S), not small (30S), ribosomal subunit. Indeed, RNA gel blot analysis showed that, under only cold stress, the accumulation of the matured 23S rRNA was greatly reduced in the *tsv3* mutant, but nearly reached WT level at high temperatures (Supplemental Figure 3). Interestingly, only under cold stress, the *TSV3* affected the 50S ribosome assembly, in turn, produced the albino phenotype. Accordingly, the loss of *TSV3*-mediated *Rpl21-23S rRNA* mRNA regulation might produce thermo-sensitive virescent phenotype under cold stress.

Nevertheless, irrespective of low or high temperatures, another *TSV3* homolog, *OsObgC1* (*LOC_Os07g47300*), null mutants (*obgc1-d1* and *obgc1-t*) and *OsObgC1* knockdown (*obgc1-d2*) mutant in rice were reported to severe or partially chlorotic phenotype during early leaf development like *AtObgC* RNAi *Arabidopsis* lines exhibiting chlorotic phenotypes (Bang et al. 2012). Additionally, the mutation of *OsObgC1* has little effect on the expression of other rice Obg homologs such as *OsObgC2/TSV3* and *OsObgM* (Bang et al. 2009). In addition to its structural similarity to both OsObgM and AtObgM in mitochondria, the obvious differences both in OCT regions (Supplemental Figure 2) and in sub-branch (Figure 4B) between TSV3 and AtObgC1/OsObgC1 strongly supported the existence of the difference in response to environment changes (e.g. temperature/light) and regulating pathways. The findings highlighted the notion that even highly conserved genes within the same or across species might play diverse and complex roles than previously recognized. Also, our observations provided the evidence of versatile roles for plant ObgCs in development. Taken together, *TSV3* is required for chloroplast development during the early leaf stage under cold stress.

### *TSV3* may be important for recovery from cold stress

In this study, the *tsv3* mutant is a typical thermo-sensitive virescent mutant, similar to the previously described thermo-sensitive virescent /albino rice mutants such as *v1* (Kusumi et al. 1997), *v2* (Sugimoto et al. 2004 and 2007), *v3* and *st1* (Yoo et al. 2009), *wlp1* (Song et al. 2014), *osv4* (Gong et al. 2014) and *tcd9* (Jiang et al., 2014). Despite the similar phenotypes in *v1, v2, v3, st1, wlp1, osv4* and *tcd9* mutants, the mechanisms and regulations affecting the chloroplast development are possibly different. Briefly, the *V1(NUS1)* encoding chloroplast-localized protein NUS1 regulates ribosomal RNA transcription under low temperature (Kusumi et al. 2011) and the *v1* mutation severely blocks the accumulation of PEP subunits (Kusumi et al. 2004). The *V2*, which encodes plastid/mitochondrial guanylate kinase (pt/mt GK), regulates guanine nucleotide pools (Sugimoto et al. 2007) under low temperature and the *v2* mutation blocks the formation of functional PEP (Sugimoto et al. 2004). The *V3* and *St1*, encoding the large and small subunits of ribonucleotide reductase (RNR) respectively, are required for chloroplast biogenesis during early leaf development and the *v3* and *st1* mutants withered to death at approximately 30 d after germination under 20°C conditions (Yoo et al. 2009). In *wlp1* mutant, the mutation of the rice large subunit protein L13 lead to abnormal chloroplast development under only cold stress (Song et al. 2014). Additionally, the mutation of rice *TCD9*, encoding α subunit of chaperonin protein 60 (Cpn60α), hinders *FtsZ* transcription/translation, in turn, influences plastid division and finally leads to abnormal chloroplasts under cold stress (Jiang et al. 2014). The mutation of *OsV4*, encoding a novel chloroplast-targeted PPR protein, leads the dramatically reduced transcriptions of some ribosomal components and PEP-dependent genes under cold stress (Gong et al. 2014).

Obviously, the loss of *TSV3* function produced low-temperature virescent phenotype before 4-leaf stage, indicating that *TSV3* was involved in a pathway that may be only required under cold stress. This was strongly supported by the highly expressed level at 20°C, regardless of WT or mutant (Figure 3B). It has been reported that cold stress interfered with protein biosynthesis in plastids by delaying translational elongation (Grennan and Ort 2007), and virescence/thermo-sensitivity played a role in protection from photo-oxidative damage before healthy chloroplasts were developed (Zhou et al. 2009). Previous reports also confirmed that the deficiency of plastid translation often led to a cold sensitive phenotype (Tokuhisa et al. 1998; Ahlert et al. 2003; Rogalski et al. 2008; Liu et al. 2010). Taken together, *TSV3* might be involved in a protection mechanism under cold stress and the reduction of TSV3 would lead to a cold-sensitive chloroplast deficiency.

In conclusion, our data clearly indicated that *TSV3* was fundamentally involved in the biogenesis of plastid ribosomes under cold stress during chloroplast development in early leaves and a hypothetical model for *TSV3* function was shown in Figure 9. In this model, under cold-induced conditions, TSV3 was translocated into chloroplast to interact with the chloroplast ribosome 50S to produce active PEP capable of transcribing photosynthetic or some housekeeping genes, unlike *AtObgC/OsObgC1* which is translocated into chloroplast under light-induced conditions (Bang et al. 2012). It merits further investigation in the extent of *TSV3* function variation, which might control expression of active PEP, according to cell type, developmental stage or environmental conditions.

**Figure 9.**
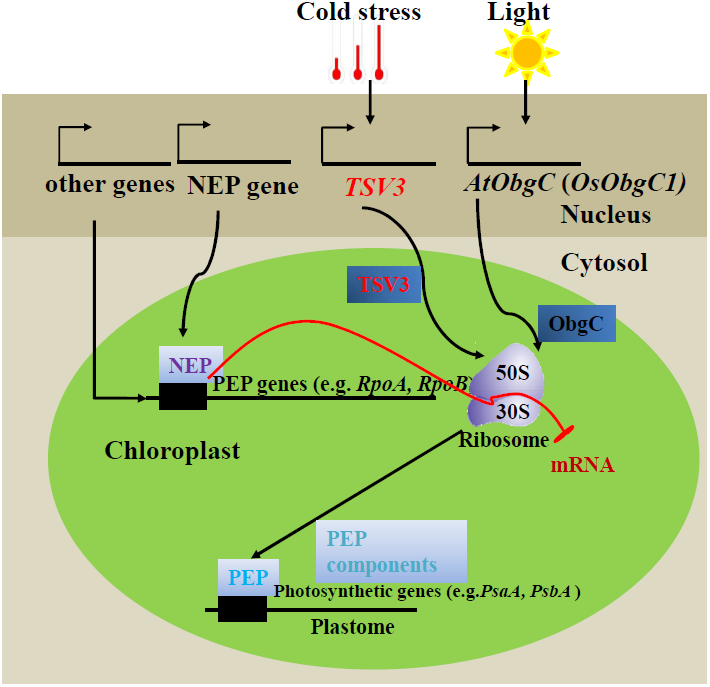
A functional model of *AtObgC* in *Arabidopsis* (Bang et al.2012) and the possible model of *TSV3* in developing chloroplast. Two types of RNA polymerases (NEP and PEP) have been identified in higher plant chloroplasts. TSV3 interacts with the rice chloroplast 50S ribosome subunit, which functions in the translation of protein encoded by the chloroplast gene and also regulates the transcription of photosynthetic or some housekeeping genes by impacting PEP synthesis. In the absence of TSV3, PEP activity stays low, which couples biosynthesis of chlorophyll and proteins, is significantly reduced at the seedling stage under cold stress, leading to the virescent phenotype.

## ACKNOWLEDGMENTS

We sincerely thank Dr. Youbin Xiang (Stowers Institute for Medical Research, USA) for her critical reading of and suggestions for our manuscript. We are grateful to Prof. Zhongnan Yang for kindly providing pMON530-GFP vector and for his constructive comments and suggestions on this paper as well. The project was financially supported by the Natural Science Foundation of China (No. 30971552), Minister of Science and Technology of China (MOST) (2016YFD0100902), the Shanghai Municipal Science and Technology Commission (16ZR14253000 and 16391900700), and Innovation Program of Shanghai Municipal Education Commission (2017-01-07-00-02-E00039).

---

**Supplemental Figure 1.**
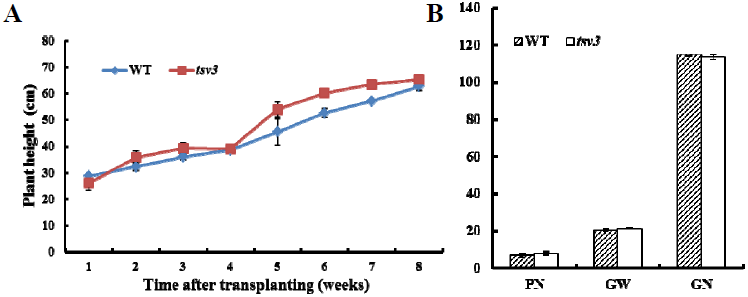
Characters of the *tsv3* mutants and WT plants grown in fields (2010, Shanghai, China). A Changes of plant height from transplanting to heading. B Comparison of three yield-related traits between WT and *tsv3* mutant. WT, wide type; PN, panicle number per plant; GW: 1000-grain weight (g); GN, grains per panicle

**Supplemental Figure 2.**
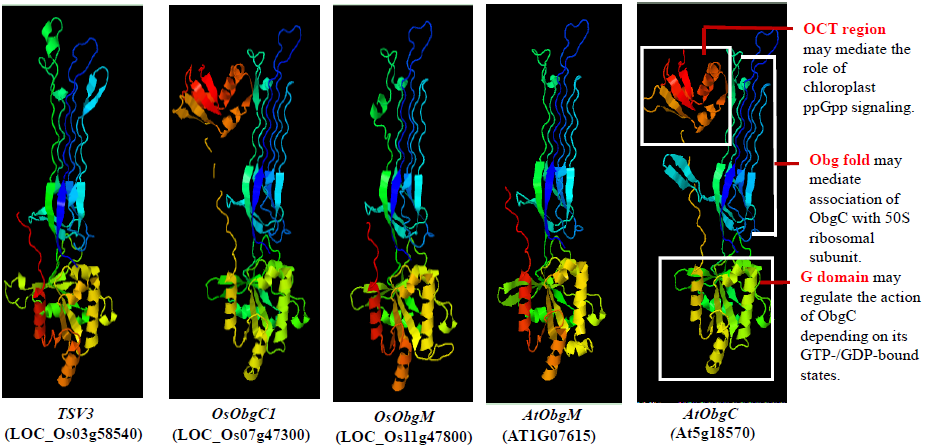
Predicted 3D structure of TSV3 and homologous proteins. Data predicted by using the Phyre 2 server (http://www.sbg.bio.ic.ac.uk/phyre2/html/page.cgi?id=index) and functional mode of three Obgc domains in AtObgC were cited from Bang et al.(2012).

**Supplemental Figure 3.**
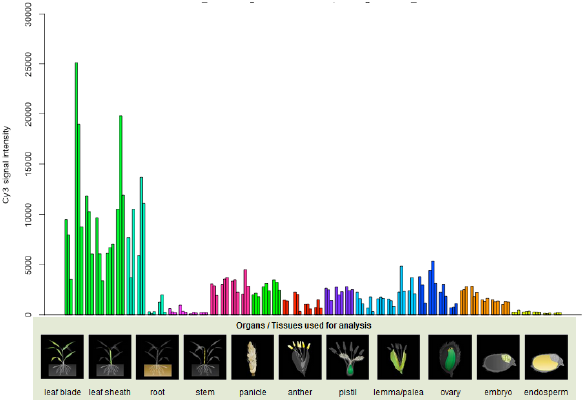
Expression patterns of *TSV3* (*LOC_Os03g58540*). Data were cited from the rice expression profile database, RiceXPro (http://ricexpro.dna.affrc.go.jp/category-select.php).

**Supplemental Figure 4.**
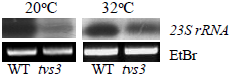
RNA blot analysis of accumulation of chloroplast 23S ribosomal RNA. Five micrograms of total RNA from wild-type and tsv3 3-leaf-stage seedlings grown at 20°C and 32 °C was extracted. RNA gel blot analysis was performed as described previously (Chi et al., 2014, published in The Plant Cell, Vol. 26: 4918-4932). In addition, the 25S rRNA stained with ethidium bromide (EtBr) is shown as a loading control.

## Table legends

**Supplemental Table 1** The PCR-based molecular markers designed for fine mapping

**Supplemental Table 2** Markers designed for realtime RT-PCR

